# SANTA-SIM: Simulating Viral Sequence Evolution Dynamics Under Selection and Recombination

**DOI:** 10.1101/348508

**Authors:** Abbas Jariani, Christopher Warth, Koen Deforche, Pieter Libin, Alexei J Drummond, Andrew Rambaut, Frederick A Matsen, Kristof Theys

## Abstract

Simulations are widely used to provide expectations and predictive distributions under known conditions against which to compare empirical data. Such simulations are also invaluable for testing and comparing the behavior and power of inference methods. We describe SANTA-SIM, a software package to simulate the evolution of a population of gene sequences forwards through time. It models the underlying biological processes as discrete components: replication, recombination, point mutations, insertion-deletions, and selection under various fitness models and population size dynamics. The software is designed to be intuitive to work with for a wide range of users and executable in a cross-platform manner.

## 1. Introduction

Simulating population dynamics is a popular and effective strategy to model the outcome of molecular genetic processes (e.g., selection and recombination) and to verify evolutionary hypotheses against experimental observations. Simulations of evolutionary histories in population genetics can be categorized either as forward-in-time or backwards-in-time (coalescent) genealogical models. Coalescent models have been historically the leading simulation method and are used for the inference of genetic variation in populations through progressively coalescing lineages according to a stochastic process until only the most recent common ancestor of the sample population is reached [1, 2]. This process is appreciated for its time and memory efficiency as it only considers a sample of observed individuals, irrespective of how large the population is, and is widely used to simulate changes in population size, recombination and sub-populations with migration [3, 4]. By contrast, forward-in-time evolution simulations are computationally more intensive as the evolutionary history of the entire population is modelled through time. However, these models allow for more complexity and scenarios can include population processes (e.g., natural selection) that are difficult to incorporate in the backwards-time approach [5]. A large collection of software tools for genetic data simulation has been developed in the past decades. Extensive descriptions of available forward simulators show that a wide but incomplete range of genetic and population processes are considered by these tools [6, 7, 8, 9].

We present the SANTA-SIM software package, which implements an individual-based discrete-generation forwards-time simulator for molecular evolution in a finite population. SANTA-SIM is directed towards haploid organisms, and particularly useful to study rapidly evolving pathogens such as RNA viruses that can experience diverse selection pressures and recombination events. SANTA-SIM is distinguished from previous simulators (of virus evolution) by its modularity, flexibility and extensibility of simulation components. Discrete components reflect the different underlying biological processes, which can be configured separately and combined to simulate complex evolutionary processes. Unlike many of previously published tools designed to study population genetics [10, 11, 12, 13, 14, 15], SANTA-SIM is focused more on the dynamics of selection in the face of a changing fitness landscape. For instance a simulation could have consecutive epochs to model environmental changes. Moreover, rather than only modelling selection by predefined fitness values for different alleles, fitness of each allele could be affected by context dependent effects such as the population size or the duration that the allele has been present in the population. Samplers can be adapted to extract statistics, sequence alignments and genealogy trees from the simulation. Furthermore, SANTA-SIM allows simulation of insertion-deletion mutations (indels) and dynamics of population size. Despite recent advances in simulation models [16, 17], to our knowledge, none of the previously published simulators are able to sample genealogy trees, simulate indels and such complex selection scenarios concurrently.

## 2. Software features

SANTA-SIM was written in the Java programming language and is available as an open-source project (https://github.com/santa-dev/santa-sim) under the Apache License Version 2.0 (APLv2). A simulation run only requires a pre-built cross-platform executable file and an XML configuration file, where the configurable aspects of the simulation process are detailed. The SANTA-SIM framework was developed in a modular manner so that different biological processes are implemented in separate components. This design allows users with basic programming experience to understand how the software functions, and to easily adapt existing or implement new features.

An overview of the evolution simulation process is demonstrated in Figure 1 for a cycle of two consecutive generations. After the calculation of fitness for the individuals within a population, the next generation is selected from these parents according to their fitness. Recombination can occur between two parents to generate a progeny with a genome inherited from two different parents. Subsequently, mutations are introduced into the new generation and the simulation will proceed for the following generations.

**Figure 1:**
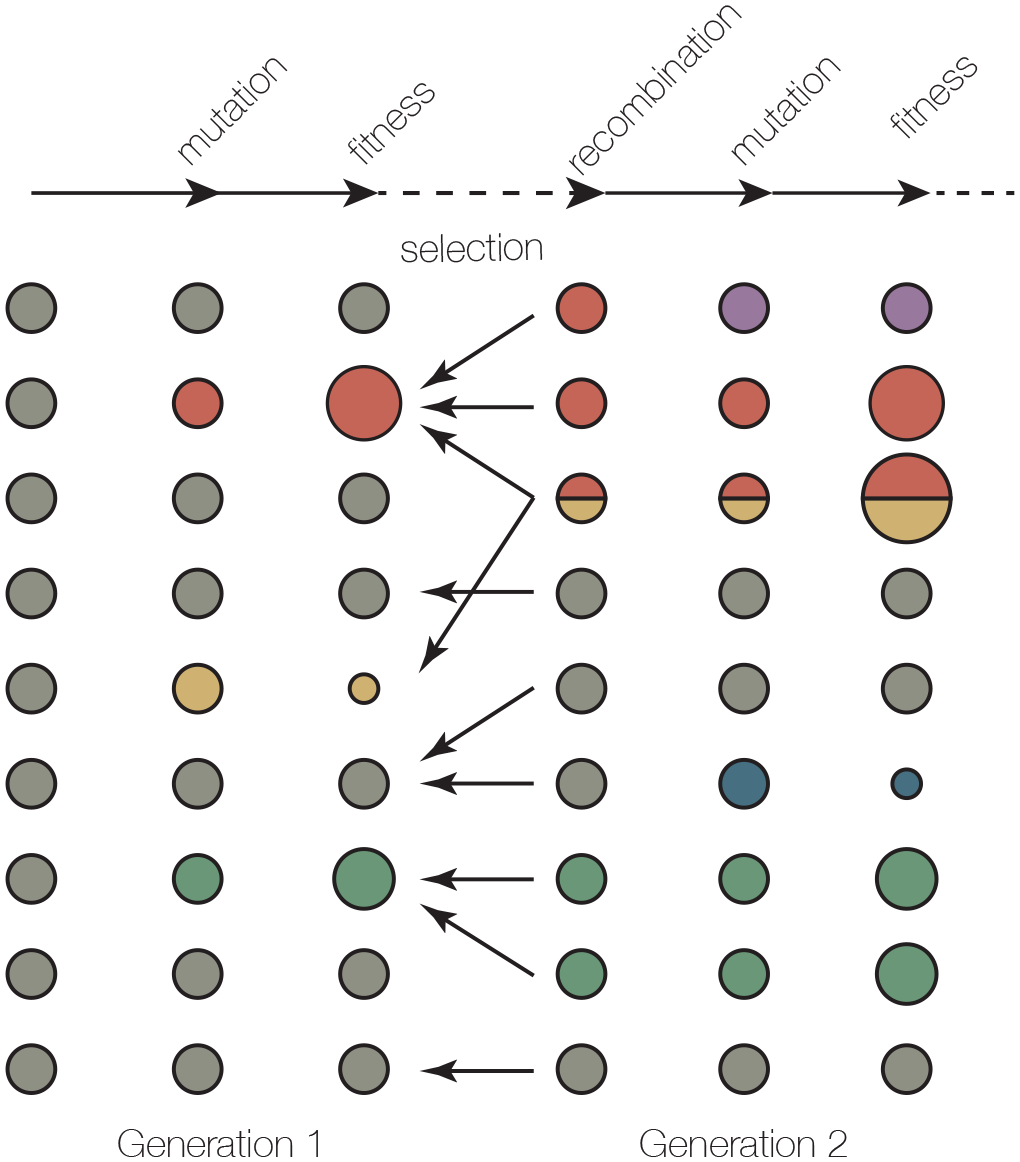
Overview of simulation process in SANTA-SIM. A cycle of two generations in SANTA-SIM simulation, consisting of mutation, recombination, fitness evaluation and selection. The circles on the left and right respectively represent the individuals from the first population (parents) and the second generation (progenies). The size of the circles represent the fitness while the color represent the genotype. Parents with higher fitness are more likely to be selected to generate a progeny, shown by the number of arrows. Each progeny could be generated from one parent (clonal replication) or two parents (recombinant replication).

In order to enable SANTA-SIM to be easily extended, the most fundamental simulation components are presented to the developer as Java interfaces. An overview of the interfaces, and their default implementations within SANTA-SIM can be found in Table 1.

**Table 1:** An overview of the Java interfaces that correspond to the basic simulation components

| **Interface** | **Component description** |
| --- | --- |
| FitnessFactor | Enables developers to define a custom fitness function. Default implementations of this interface implement fitness functions that take into account genome sequence and the size of the virus population. |
| Mutator | Enables developers to control how a molecular sequence is to be modified when a mutational event takes place. SANTA-SIM ships with a default implementation that allows mutations to happen on the nucleotide level. |
| Replicator | Enables developers to control how a virus can be replicated. Default implementations of this interface implement clonal replication, recombinant replication and recombinant replication with hotspots. |
| Sampler | Enables developers to implement different ways to sample from the virus population. Default implementations of this interface implement an alignment sampler, an allele frequency sampler, a genome description sampler a general statistics sampler and a tree sampler. |
| Selector | Enables developers to implement different ways to apply evolutionary selection on individuals. SANTA-SIM ships with a binary search selector and a roulette wheel selector for constant population size. For simulating the dynamics of population size under logistic growth model a selector is implemented that takes into account the growth rate, carrying population and population size to determine the expected number of progeny for each individual. |
| PopulationGrowth | Facilitates further development of the software to specify the growth processes custom-tailored for specific cases which are not predicted in the selector interface implementations. |

Using these model components, the entire simulation can be organized as a sequence of epochs, with each epoch having different selection functions, replication operators and mutation operators. The transition of the population to a new epoch thus reflects a deterministic environmental change for the population. The software can also be configured to report insights into the operation of the simulation. These outputs can either be at the population level by describing allele frequencies through time, population fitness, diversity and divergence or at the individual level by providing nucleotide or amino acid sequence alignments at chosen loci and phylogenies.

### Population, individuals, genomes and features

The population in SANTA-SIM consists of individual organisms, each of which contains a single genome. The genome is a linear sequence of nucleotides organized into *features*. Features reflect the genome organization into genes and open reading frames. Each feature may be composed of one or more fragments of the genome, read in either a forward or reverse direction, and may overlap with each other. Different modes and degrees of selection on either nucleotides or amino acid sites can be specified on each feature.

### Evolutionary process

The evolutionary process in SANTA-SIM assumes discrete generations, and each generation consists of a concatenation of discrete components. The population is subjected to mutations, recombination and various models for fitness assignment based on the genotype of the genome features. The size of the population can change depending on the overall fitness of the population. The interaction between these components supports the configuration of complex evolutionary scenarios.

The simulation begins with an initial population of individuals which is seeded from a single sequence or a pool of different sequences. At each generation, evolution is simulated in four sequential steps of fitness, selection, replication and mutation.

### Fitness calculation

The fitness of each genome is calculated using one or more fitness functions (Table 2). By shuffling selection coefficients among states over time, non-stationary random positive selection can be implemented. Distinct fitness functions can be defined for the nucleotide sequence and the corresponding amino acid translation.

**Table 2:**
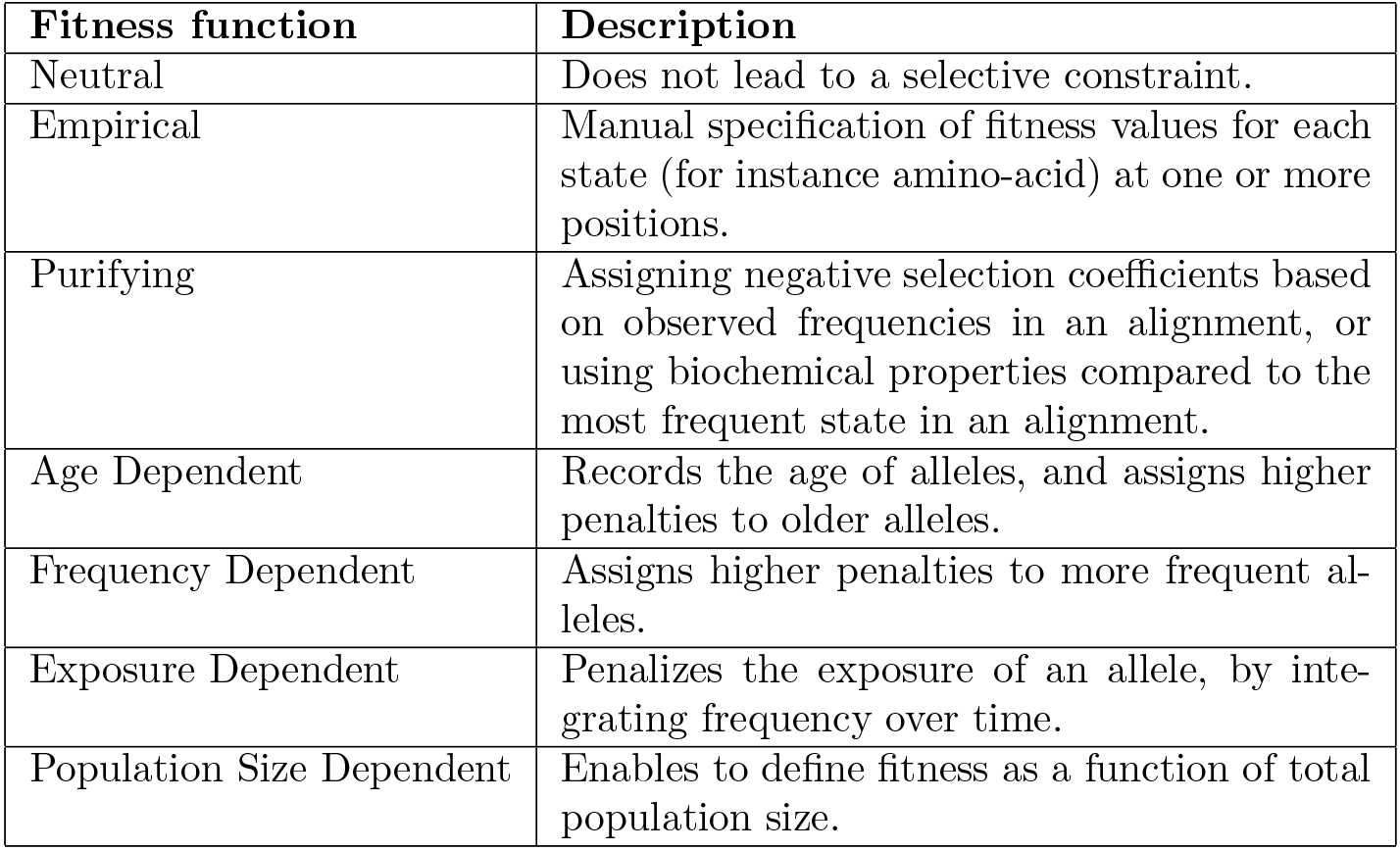
Overview of the fitness functions that can be implemented in the simulations by their respective specification in the XML file.

Furthermore, different regions of the genome can be assigned different fitness functions (e.g., most sites under purifying selection but with a few sites under diversifying selection). Such fitness functions are introduced in the XML configuration file.

### Selection

The next generation of individuals then selects their parents from the previous generation where each parent is selected with replacement with a probability proportional to its genome’s fitness. The number of parents that are selected for each new individual depends on the mode of replication, which is described next.

### Replication

Together with the mutation component, below, the replication component is analogous to the actions of a polymerase complex and produces the genetic material for a new individual from one or more parents. The simplest replicator is clonal and the descendant inherits the genome of exactly one parent. We also provide a recombinant replicator that models a ‘template-switching’ polymerase. For this replicator, two probabilities are defined: a probability that a recombinant is used instead of the clonal replicator, and a probability of the polymerase switching between the parents’ templates as replication proceeds along the genome.

### Mutation

Mutation is implemented as an independent process after replication. The user specifies a per-site and per-generation probability of mutation, and the mutator component then applies mutations to the genome accordingly. For efficiency, the default mutator draws the number of mutations from a Poisson distribution with an expectation given by the number of nucleotides and the mutation rate. These are then distributed uniformly across the sites. A bias towards transition-type mutations can be specified to reflect the action of specific polymerases.

> In addition to substitution mutations, SANTA-SIM also offers an indel mutation model that can be useful to more closely mimic the evolutionary behaviour of retroviruses like HIV-1. The user can specify a per-generation probability of insertion or deletion, together with an independent distribution of indel length. Frame-shifting a genome is assumed to be nearly always fatal so only whole-codon indels are permitted. Indels not only change the content and length of a simulated genome, but they can also affect regions subject to fitness constraints. Depending on the position relative to the boundaries of fitness-constrained regions, indels can either shrink or lengthen a feature and therefore have an impact in fitness scores. Changes to fitness-constrained regions will affect subsequent generations of the lineage, but will not affect sibling lineages. Details on the indel mutation model and illustrations of the implications for fitness calculations can be found at the project webpage.

### Sampling sequences, phylogenies and statistics

Given the large scale of many typical simulations with biologically relevant parameters, we have made every effort to use memory efficiently. For example, genome sequences are stored in a central ‘gene-pool’ so that only unique genomes are stored with the individuals having only an index for the genome they currently carry. Individuals that replicate without any mutations thus inherit this index. This also makes calculations of the population genetic diversity more efficient. In addition, where applicable, fitness may be computed incrementally from the parent’s fitness in the previous generations and incidental mutations. We also implemented an optional framework where genomes are stored as differences from a central ‘master’ sequence. This master sequence can be recalculated occasionally to release memory.

At predefined time intervals or specific times during the simulation, SANTA-SIM can report statistics about the current population, including average fitness, genetic diversity and number of unique genomes. A random sample of individuals of a specified size can also be generated and the genomic sequences recorded as a nucleotide or amino acid alignment (FASTA or NEXUS format) for use in other software applications.

Finally, it is possible for SANTA-SIM to keep track of the genealogy of the entire population and then provide the tree of the individuals sampled. A variety of events can be recorded, such as the prevalence of different states in selected sites, or shuffling events in the purifying fitness function, to be able to investigate the effect of these events on the population.

## 3. Simulation examples

We describe three example runs that demonstrate some of the functionalities of SANTA-SIM and the fitness functions that can be applied. A first example simulates the consecutive selection of a set of deleterious mutations that each give a fitness advantage in a new environment. A second example demonstrates how the frequency of initial beneficial mutation during a selective bottleneck impacts diversity and phylogeny, which is related to the discussion of soft and hard selective sweeps [18]. A third example shows the interplay between the fitness of a mutation and its frequency, as in the case of a host-pathogen interaction. The configuration files and additional details of these simulations are available as supplementary material.

### 3.1. Directional selection driven by changing selective advantage of mutations

In the face of environmental changes, certain mutations which were previously neutral or deleterious can confer a selective advantage in the new environment.

Such an scenario occurs when a viral population, e.g. HIV-1, is subjected to suppressive action of antiviral treatment and demonstrates adaptive evolution. We can simulate this example using two epochs, with no selective pressure present in the first epoch. A population of 10,000 sequences was created by evolving an initial sequence (609 nucleotides in length) under mutation and moderate level of purifying selection for a duration of 3,000 generations. A set of four mutations across this sequence were associated with a selective disadvantage, making their fixation in the population unlikely. The second epoch started after generation 3,000, with a strong beneficial impact of these four mutations in the new environment, and their subsequent selection. Figure 2 illustrates the dynamics of population diversity and the frequency of the beneficial alleles while Figure 3 shows the phylogeny reconstructed from sequences sampled through the course of the simulation. Genetic diversity was calculated as the mean pairwise distance, based on percent identity, within a random sample of 1000 individuals from the population.

**Figure 2:**
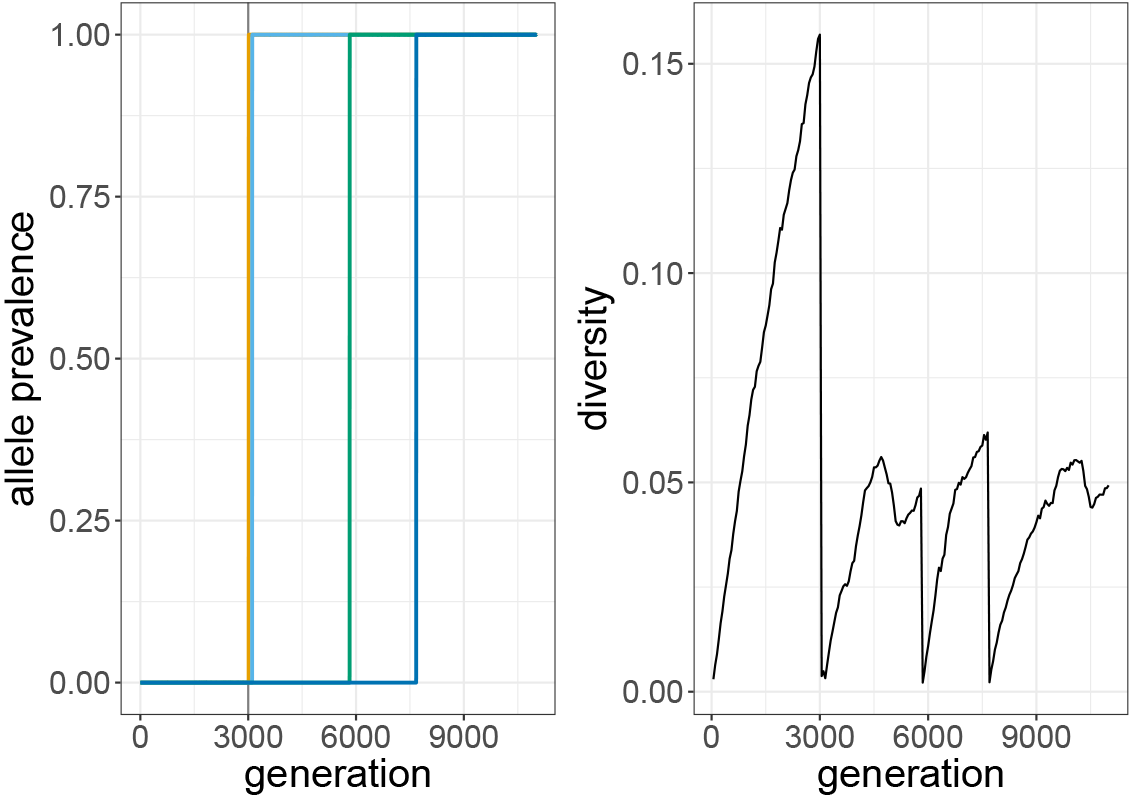
Diversity and allele prevalence through several selective sweeps. The simulation has two phases. In the first 3000 generations, the only selective force is purifying selection. After this initial phase (gray line), 4 particular mutations become beneficial:50T (yellow), 100K (light blue), 150A (green) and 200G (dark blue). Mutation 100K has been present in the initial population in low prevalence. Diversity drops through each wave of selective sweep where a beneficial mutation appears and takes over. The simulation starts from a population with only one sequence at the first generation. Diversity was defined as the mean pairwise identity percentage between all sequences. For a given nucleotide position between two sequences, two non-identical bases will result in a zero score for that position while identical bases give a score of one, The distance of the two sequences was calculated as the mean of such identity scores across all nucleotide positions.

**Figure 3:**
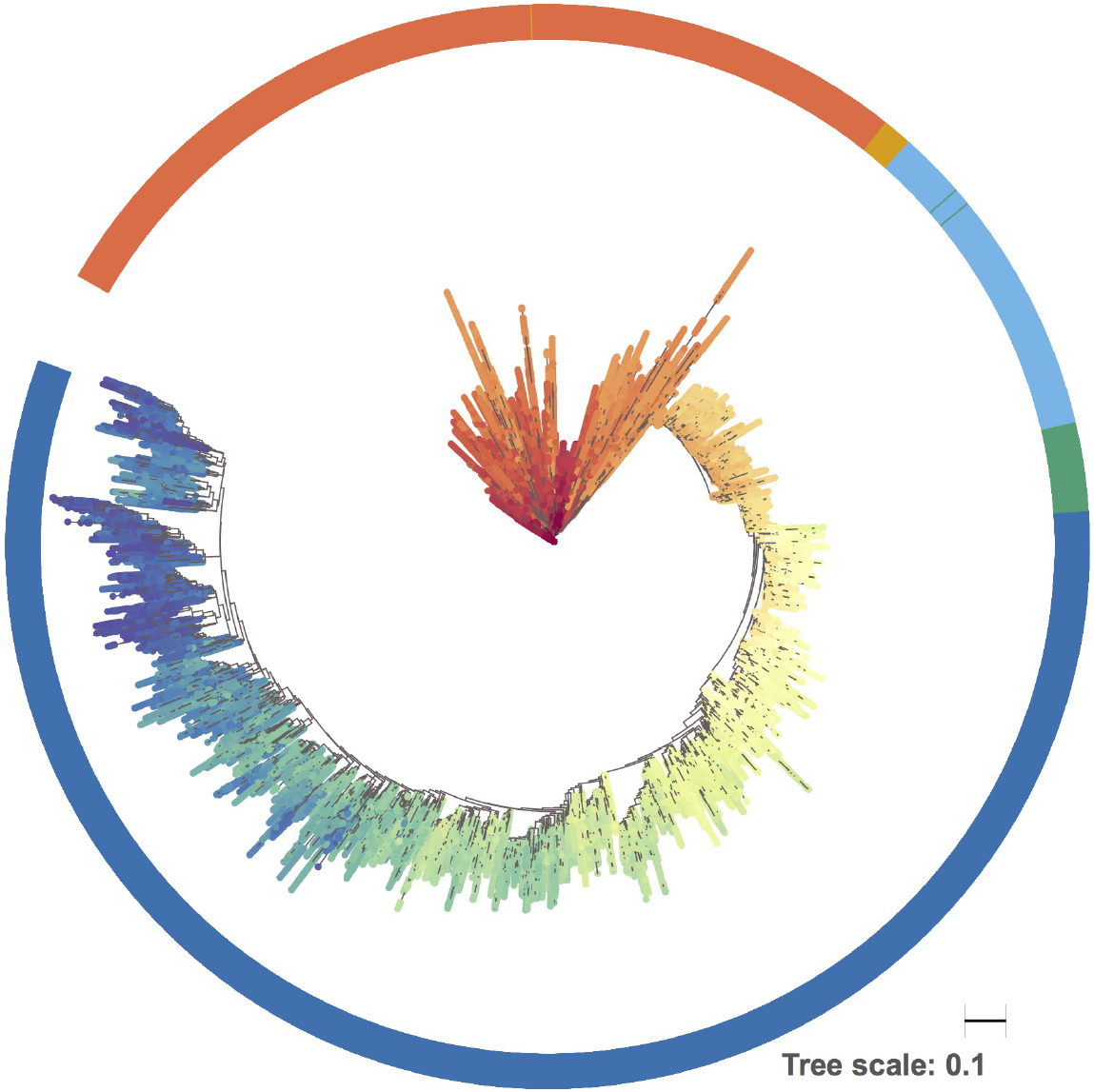
Phylogenetic tree from the sampled sequences through multiple waves of selective sweep. From the simulation run of 10,000 generations, a sample of 500 sequences was collected at every 100th generation, and a tree was made with FastTree. The tree is coloured by increasing generations (from red to blue) and the outer band denotes the consecutive selection of beneficial mutations (see Figure 2 for mutation colors).

### 3.2. Fraction of individuals with beneficial alleles affects the trajectory of a selective sweep

Directional selection, as shown in the previous section, results in a decrease of genetic variation in the viral population, but the extent of reduction depends on the initial frequency of the beneficial mutation and the strength of selection [19]. When a strongly beneficial but rare mutation increases in frequency, the genetic background of this adaptive mutation will dominate the population and have a strong impact on diversity. In contrast, rapid adaptation to a novel selection force by a mutation either highly prevalent or newly arising simultaneously in different individuals will lead to a less drastic reduction in genetic variation of the population [20]. This example investigates how an initial mutation frequency affects a selective sweep. Two independent simulations were carried out for a starting population of size 10,000, but where one population had 55 individuals with the beneficial mutation before the onset of the selective sweep, while the other population only had two individuals with the beneficial mutation.

Figure 4 illustrates that the magnitude of diversity reduction upon a sweep is dependent on the fraction of the beneficial mutation in the initial population. Diversity remains at a quasi-constant level after the sweep since the span of the number of generations here is rather short.

**Figure 4:**
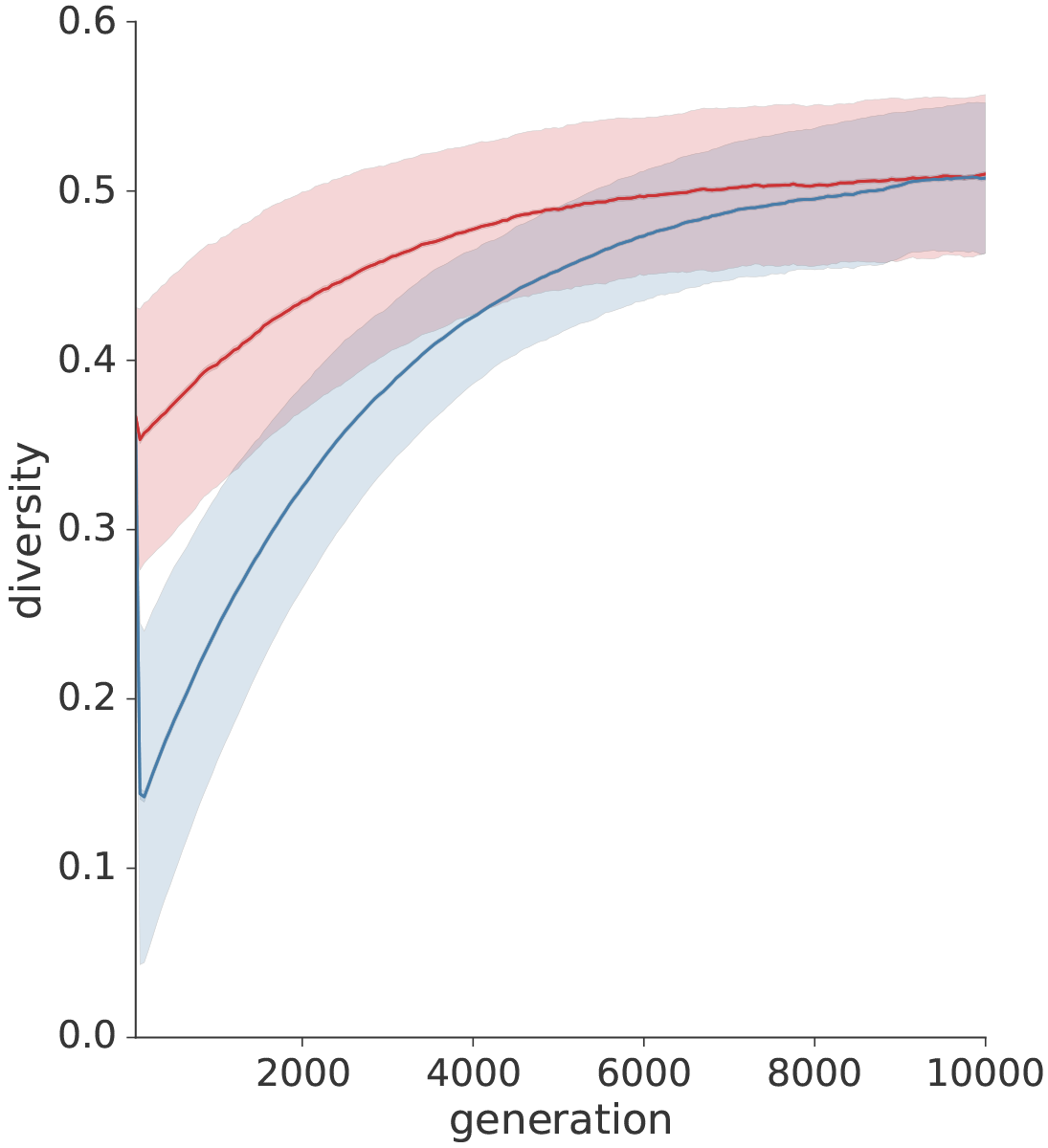
Diversity trajectory through selective sweeps for different initial frequencies of the beneficial alleles. Two modes of selective sweep are simulated for a population of size 10,000: In one case a small fraction of the initial population carries the beneficial allele (two in 10,000; blue line) while in the other case a higher fraction carries this allele (55 in 10,000; red line). The diversity is calculated as the mean pairwise distance between 1000 sampled sequences from the population, similar to the previous simulation. The simulation is repeated 1000 times. The mean and standard deviation of these replicates are shown shown as the thick line and shaded area respectively.

In Figure 5 phylogenetic trees constructed after fixation of the beneficial allele demonstrate that the branching pattern at the root of the trees is distinguishably different for the two modes of the selective sweep, which resulted in the pattern in Figure 4.

**Figure 5:**
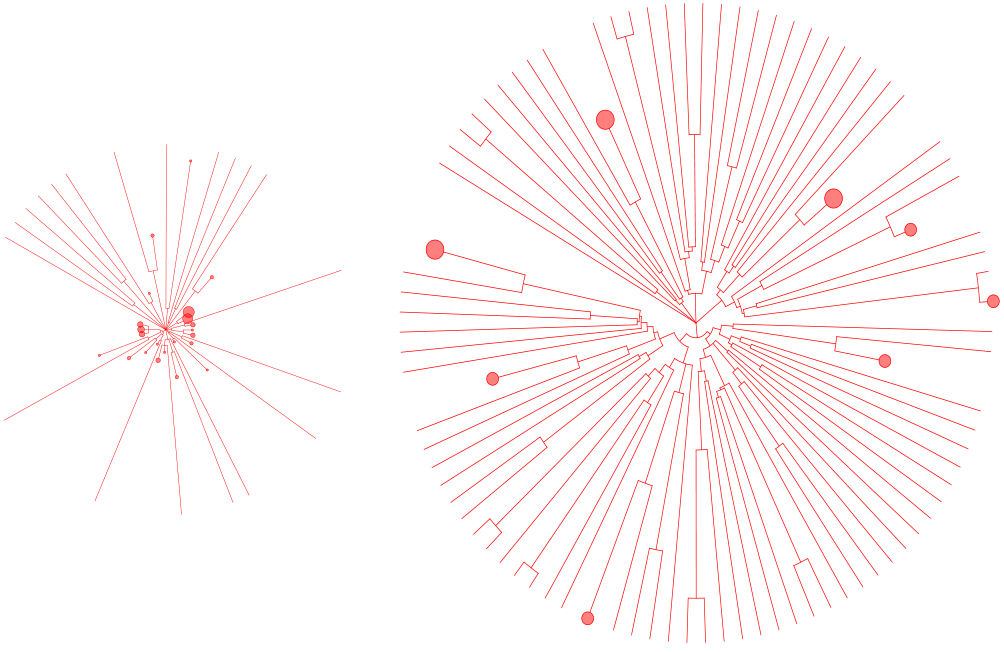
Phylogenetic trees after fixation of the beneficial allele grouped for two levels of initial prevalence of allele. A population with two levels of initial prevalence of a beneficial allele is subjected to selective sweeps. Phylogenetic unrooted trees are sampled (100 sequences), using tree sampling capability of SANTA-SIM, after the fixation of the beneficial allele. The tree on the left corresponds to the case where the starting population had a lower number of individuals with the beneficial allele (2 in 10,000), while the tree on the right corresponds to the case where the starting population had a higher number of individuals with the beneficial allele (55 in 10,000). Clades were collapsed (red dots) when average branch length distance to the taxa were below an illustrative threshold.

### 3.3. Simulation of dynamics of host-pathogen co-evolution: A complex selection scenario

Pathogen escape from the host immune response can be transient when adaptation of the host immune system to the pathogen variant occurs. In this example, we simulated a scenario where the pathogen has the capability to develop escape from the host’s immune system and increase its fitness by acquiring a particular beneficial mutation. At the same time, the host gradually develops more protection against the alleles of the pathogen exposed to the host.

SANTA-SIM can model this kind of adaptive dynamics using an *Exposure Dependent Fitness Function* which assigns fitness values to pathogen variants based on the exposure of their allele in the population since it least appeared. Such fitness is assigned as exp(penalty_parameter ∗*E*) where *E* is the integrated prevalence of the allele over time since its last appearance. Thus the variants are penalized when they have been present for a longer period at a higher prevalence in the population. The severity of how much the exposed alleles are punished also depends on a penalty parameter. Overall, the fitness of each individual is determined by two factors: presence of the beneficial mutation and the exposure penalty which models adaptive evolution of host’s immune system.

Figure 6 demonstrates the effect of the exposure penalty parameter. A low value (shown in green) is similar to the absence of exposure-dependent fitness function and the beneficial mutation reaches and remains a high prevalence. The minor fluctuation in prevalence results from the marginal increase relative to other alleles with selection coefficient of 0.05. For higher values of the penalty parameter for allele exposure, it is observed that mutations that increase in prevalence are penalized for their exposure and affecting the duration of high prevalence. Simulation for each parameter is shown for duplicates.

**Figure 6:**
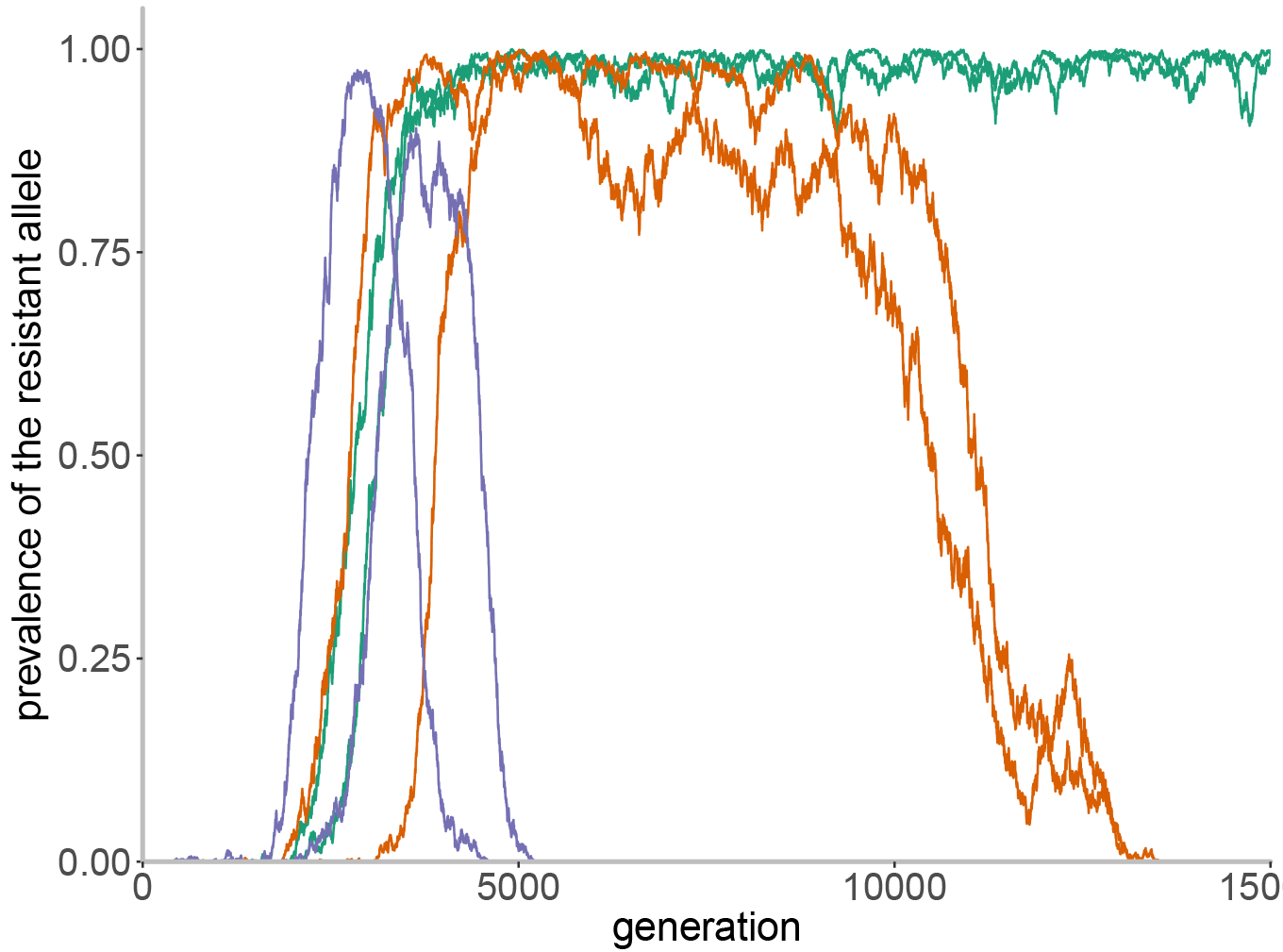
Simulation of selection dynamics in host pathogen co-evolution. Simulation of the interplay between the appearance of a escape mutation in a pathogen and host adaptation to the resistance. The selection coefficient of the beneficial mutation was set to 0.05. The exposure dependent fitness function was used to simulate the gradual decrease in fitness of the beneficial resistance allele in a pathogen as the hosts immune system adapts. Three parameters for exposure penalty were used. The penalty parameter for the green curves is set to 10^−7^, for the orange curves it is 10^−6^ and for the purple curves it is 10^−5^. There are two replicates shown for each simulation.

## 4. Benchmarking

Multiple simulation configurations were defined with a range of population sizes and genome lengths. A memory footprint and elapsed wall-clock time were measured for each simulation as shown in Figure 7. A computer with 3.4 GHz Intel^®^ Core i7 CPU and 32 GB of RAM was used to run the simulations (Ubuntu 16.04, Java openjdk version 1.8). All simulations are configured to run for 10,000 generations under purifying selection and without sampling the simulated sequences. More information on the analysis describing these performance tests is available at the project website.

**Figure 7:**
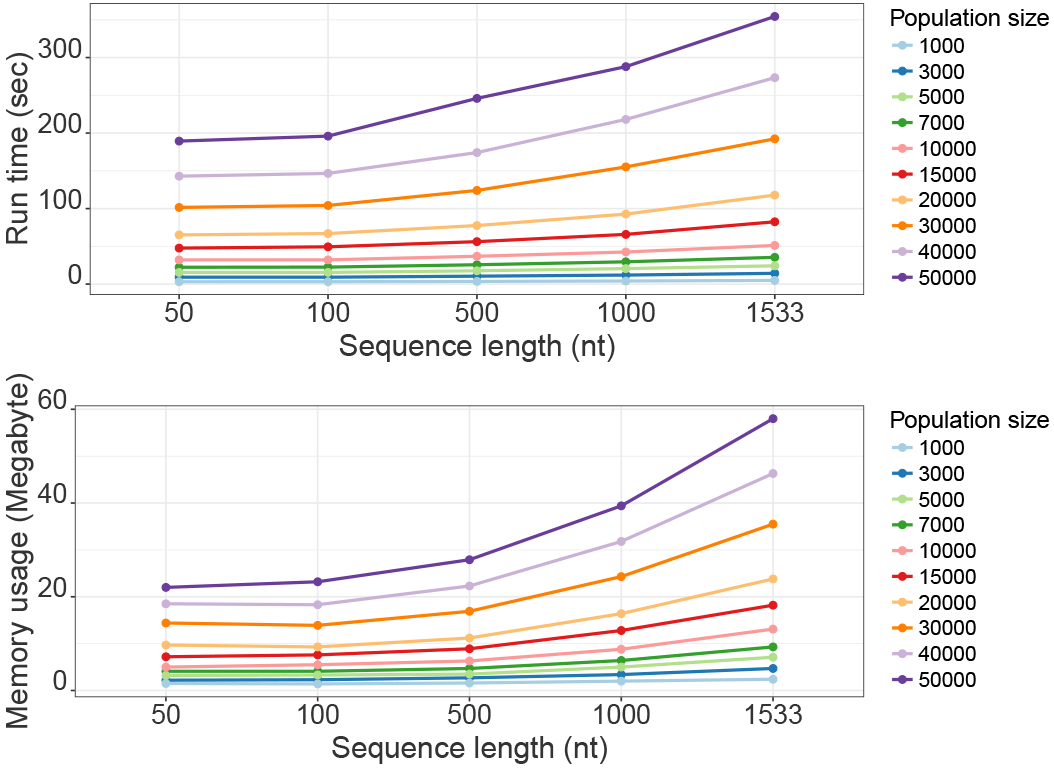
performance benchmarking. Memory and run time of simulations with purifying selection for 10000 generations with different population sizes and genome lengths.

## 5. Conclusions

SANTA-SIM is a forward-time discrete-generation gene sequence simulator, designed to scale to large population sizes and microorganism-scale genome lengths while implementing complex selection and recombination scenarios. SANTA-SIM is open-source software and written in an extremely modular fashion to facilitate a wide range of additional components being implemented to accommodate varying simulation environments and organisms.

## 6. Acknowledgments

The authors would like to thank Beth Shapiro for testing the software. CW and FM were partially supported by the Bill and Melinda Gates Foundation Award Number OPP1110049. FM was supported in part by National Institutes of Health R01-GM113246 a Faculty Scholar grant from the Howard Hughes Medical Institute and the Simons Foundation. PL was supported by a FWO PhD grant. KT was supported by a FWO postdoctoral grant. AR acknowledges the support of the Wellcome Trust through project 206298/Z/17/Z, European Union Seventh Framework Programme under Grant Agreement no. 278433-PREDEMICS and no. 725422-RESERVOIRDOCS and the Bill and Melinda Gates Foundation grant number OPP1175094.

